# Timing matters: modeling the effects of gestational cannabis exposure on social behavior and microglia in the developing amygdala

**DOI:** 10.1101/2025.02.17.638714

**Authors:** Aidan L. Pham, Ashley E. Marquardt, Kristen R. Montgomery, Karina N. Sobota, Margaret M. McCarthy, Jonathan W. VanRyzin

**Author notes:** <u>Corresponding Author:</u> Jonathan W VanRyzin, <u>Author Address:</u> Department of Psychology and Neuroscience University of North Carolina at Chapel Hill 235 East Cameron Avenue Chapel Hill, NC 27514.

## Abstract

Cannabis is the most frequently used illicit drug during pregnancy, with use steadily increasing in the United States as legalization and decriminalization expand to more states. Many pregnant individuals use cannabis to reduce adverse symptoms of pregnancy, considering it to be less harmful than other pharmaceuticals or alcohol. The primary psychoactive component of cannabis, delta-9-tetrahydrocannabinol (THC), acts on the endocannabinoid (eCB) system, yet whether it perturbs neural development of the fetus is poorly understood. Previously we have shown that androgen mediated eCB tone in the developing amygdala promotes microglial phagocytosis of newborn astrocytes which has enduring consequences on the neural circuits regulating sex differences in social behavior. Microglia are the resident immune cells of the brain and express both receptors of the eCB system, CB1R and CB2R, making them likely targets of modulation by THC. It is also plausible that exposure to THC at differing gestational timepoints can result in distinct outcomes, as is the case with alcohol exposure. To model human cannabis use during either late or early pregnancy, we exposed rodents to THC either directly during the early postnatal period via intraperitoneal (IP) injection or in utero during the prenatal period via dam IP injection respectively. Here we show that postnatal THC exposure results in sex specific changes in microglial phagocytosis during development as well as social behavior during the juvenile period. Interestingly prenatal exposure to THC resulted in inverse changes to phagocytosis and social behavior. These findings highlight the differential effects of THC exposure across gestation.

## Introduction

With decreasing perception of danger and increasing legalization of cannabis, self-reported use among pregnant populations is steadily increasing [1–3]. Pregnant individuals report using cannabis for many reasons, including to reduce adverse symptoms of pregnancy. Among pregnant individuals that use cannabis to treat at least 1 adverse symptom, most used it to treat nausea and vomiting, with the other most common reason being to reduce anxiety or for general pain [4, 5]. This wide-ranging use of cannabis leads to individuals self-reporting using at varying timepoints across their pregnancy, although on average use is reported to be higher during the first trimester than the later trimesters [1]. Because of the dose and time dependent outcomes associated with prenatal alcohol exposure [6–8], it is possible that the primary psychoactive component of cannabis, delta-9-tetrahydrocannabinol (THC), also elicits specific effects based on the timing of exposure.

THC acts on both the type 1 and type 2 cannabinoid receptors, CB1R and CB2R, of the endocannabinoid (eCB) system [9]. The eCB system of the developing brain comes online early and signaling via the endogenous ligands 2-arachidonylglycerol (2-AG) and anandamide (AEA) play a critical role in a plethora of developmental processes including the formation of proper brain circuitry and subsequent behavior [10–12]. Previous work from our lab has shown that endocannabinoid tone varies by sex in the developing amygdala and promotes microglial phagocytosis of newborn astrocytes which ultimately contributes to a sexually differentiated play circuitry [13, 14]. Here, we sought to model human THC exposure during either the third or second trimesters and determine the impact on microglia of the developing amygdala and subsequent social behavior. In rodents, the early postnatal period, specifically the day of birth through the first week of life, is considered analogous to the third trimester of human pregnancy, while gestational days 15-21 in the rat are analogous to the second trimester [15].

During these developmental windows, microglia, the brain’s resident immune cell, are integral to appropriate brain maturation and development. They are involved in many processes including synapse formation and pruning [16–18], and importantly performing phagocytosis, the act of engulfing and digesting various structures, cells, or debris in both healthy and disease states [19, 20]. These resident macrophages perform phagocytosis to not only clear cellular debris after injury [21], but also regulate a number of processes across healthy brain development including maintaining a balance of neural precursors in the developing cortex [22], maintaining homeostasis of the baseline neurogenic cascade in the hippocampus [23], and regulating the number of newborn astrocytes in the developing amygdala [13]. Microglia possess CB1R and CB2R, and thus are targets for modulation by THC exposure [24].

Here we test the hypothesis that THC exposure during two distinct developmental periods differentially alters social behavior as well as microglial activity in the developing amygdala. Behaviorally, postnatal THC exposure increases social play in males and females, while prenatal THC exposure decreased play in males only. Interestingly, both treatment paradigms impaired social recognition in both sexes. The results from a behavioral battery during development and the juvenile period revealed no effects of either THC exposure paradigm on weight gain, reflex development, or nonsocial behaviors such as open field, novel object recognition, and elevated plus maze. At the cellular level, we find that postnatal THC exposure increases the number of phagocytic microglia in the developing amygdala in females, while prenatal THC exposure decreases the number of phagocytic microglia in both sexes. Taken together, these findings reveal that the timing of THC exposure produces specific changes to social behaviors and differential effects on microglia of the developing amygdala.

## Materials and Methods

Below is a condensed version of the Materials and Methods. Detailed Materials and Methods can be found in Supplementary Materials.

### Animal studies and treatments

Adult Sprague-Dawley rats, obtained from Charles River Laboratories, were maintained on a 12:12h reverse light/dark cycle with ad libitum food and water. Animals were mated in our facility, and pregnant females were designated as gestational day 0 (GD0) on the date of sperm positivity. Pregnant females were allowed to deliver naturally with the day of birth being designated as postnatal day 0 (P0). On P0, pups were sexed and culled to no more than 14 pups per dam. Male and female pups were used in these studies, and treatment groups and sexes were balanced across litters. For prenatal THC exposure studies, all pups were cross fostered to untreated dams with equal numbers of males and females, and equal numbers of treated and control pups per dam. All animal procedures were performed in accordance with the Animal Care and Use Committee’s regulations at the University of Maryland School of Medicine.

BrdU (50mg/kg; Sigma-Aldrich Cat#B5002) was dissolved in saline and delivered intraperitoneally in a volume of 0.1mL per pup per day.

THC was acquired from the NIDA Drug Supply Program pre-dissolved in ethanol and prepared as a 1:1:18 solution of THC:Tween-80:saline (or ethanol:Tween-80:saline for vehicle control). For postnatal THC exposure pups were given an intraperitoneal (IP) injection of THC (5mg/kg) or vehicle either once daily for 7 consecutive days from P0 to P6 for behavioral studies or once daily for 4 consecutive days from P0 to P3 for histological studies. For prenatal THC exposure pregnant dams were given a subcutaneous (SC) injection of THC (5mg/kg) or vehicle once daily for 7 consecutive days from G15 to G21. These periods were chosen to model an exposure to cannabis during the third trimester and second trimester respectively [15]. Appropriate dosing of THC was determined by prior work [25].

### Immunohistochemistry, Image Acquisition, and Cell Counting

Rats were deeply anesthetized with Fatal Plus (Vortech Pharmaceuticals) and transcardially perfused with phosphate-buffered saline (PBS; 0.1M, pH 7.4) followed by 4% paraformaldehyde (PFA; 4% in PBS, pH 6.8). Brains were removed and postfixed for 24 h in 4% PFA at 4°C, then kept in 30% sucrose at 4°C until fully submerged. Coronal sections were cut at a thickness of 45μm on a cryostat (Leica CM2050S) and directly mounted onto silane-coated slides. Immunohistochemistry was performed to stain for Iba1 and BrdU using either fluorescent or DAB based staining protocols. Stereological cell counts were performed using StereoInvestigator (MBF Bioscience) on a computer interfaced with a Nikon Eclipse E600 microscope and MBF Bioscience CX9000 camera. Confocal fluorescence images were acquired using the Nikon CSU-W1 or A1 microscope equipped with 405, 488, 561, and 647 lasers, using a 20x (0.75 NA) and Nikon Elements software. Widefield fluorescence and brightfield images were captured on a Keyence BZ-X700 microscope using a 4x (0.2 NA), 20x (0.75 NA), 100x oil (1.40 NA) objective and BZ-X Viewer software. Density of microglia was determined by using Imaris.

### Developmental battery

The developmental battery was performed as previously described [26]. Behavior testing took place during the dark phase of the animal’s light/dark cycle under red light illumination. Weights were taken from P0-90, dam latency to retrieve pups was performed on P5, surface righting from P5-10, developmental locomotion from P5-15, negative geotaxis from P5-15, and wire hang from P10-15.

### Juvenile Behavioral battery

The behavioral battery was performed as previously described [26]. All behavior testing took place during the dark phase of the animal’s light/dark cycle under red light illumination. Animals were weaned on P21 and housed in same-sex, same-treatment sibling pairs. Open field was performed on P26, novel object recognition on P27, social recognition on P29, social play on P30, and elevated plus maze on P32.

### Quantification and statistical analysis

All values are shown as the mean ± SEM. Statistical analyses were performed using GraphPad Prism version 10.4.1 for Windows, GraphPad Software, www.graphpad.com. Data in the figures represent individual offspring values and means. Statistical details of experiments can be found in figure legends (tests used, exact n, p value).

## Results

### Postnatal THC exposure increases social play and decreases social recognition without impacting development or other nonsocial behaviors

We began by investigating the effects of postnatal THC exposure on typical developmental milestones and behavior. We treated rat pups with THC (5 mg/kg) beginning on postnatal day 0 (P0; birth) until P6 and then assessed behavior using a series of tests (Figure 1A). Overall, postnatal exposure to THC did not alter developmental locomotion, negative geotaxis reflex, surface righting reflex, wire hang ability, or body weight gain (Figure 1B-C, E-G). However, after postnatal THC exposure dams were slower to retrieve males, thereby abrogating the propensity of dams to retrieve males first [27] (Figure 1D). We then performed a series of behavioral tests during the juvenile period on these same animals (Figure 1H) and found no effect on open field center time or line crosses, novel object recognition, or open arm time in the elevated plus maze (Figure 1I-L).

**Figure 1:**
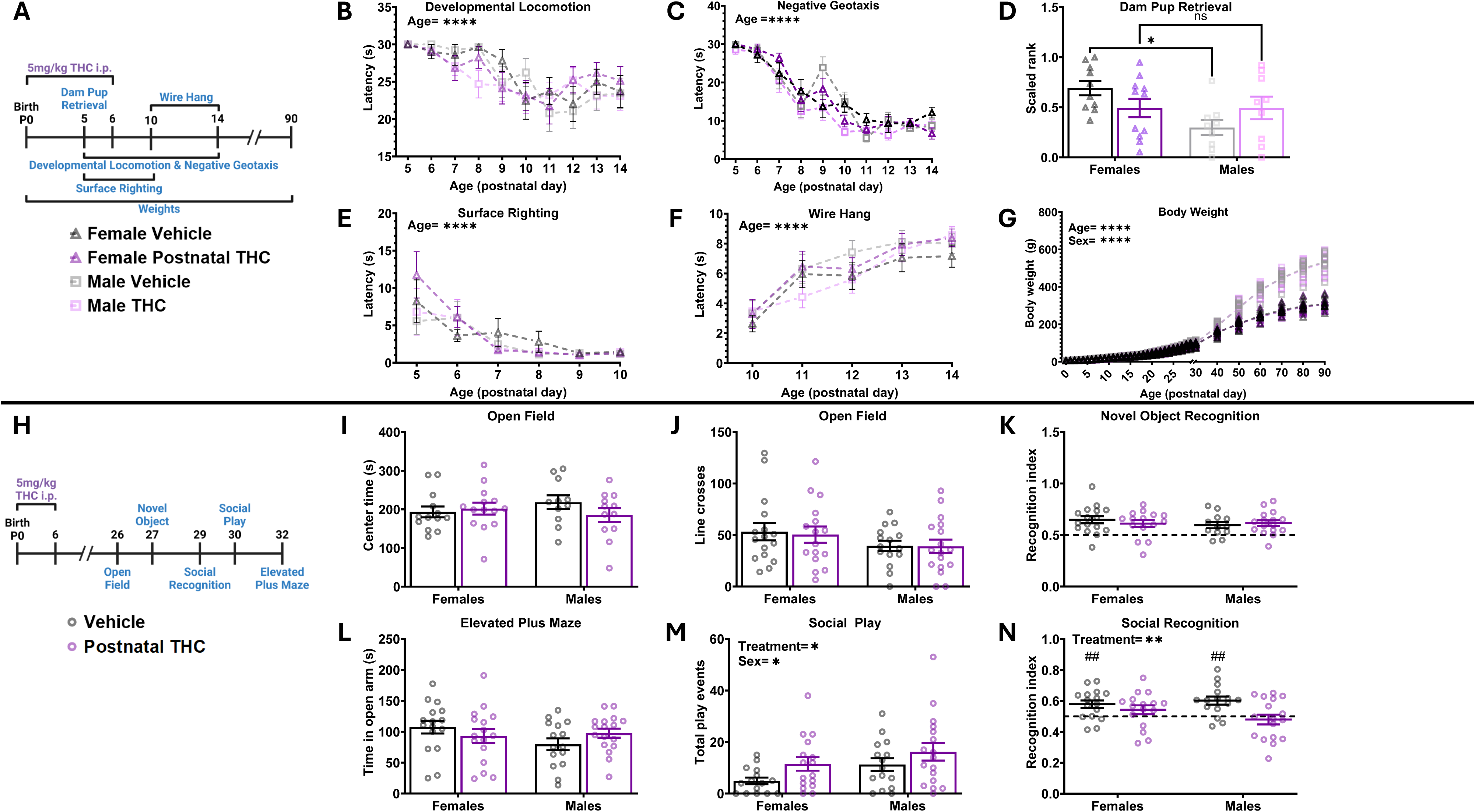
Postnatal THC exposure increases social play and decreases social recognition but does not broadly effect development milestones or other nonsocial behaviors. Bars represent the mean ± SEM, open circles represent individual data points for each animal. A. Schematic showing treatment paradigm, timeline, and legend for B-G. B. Quantification of the latency to locomote. Repeated measures three-way ANOVA revealed no effect of sex, data was then collapsed. Repeated measures two-way ANOVA showed decrease in latency to locomote as animals aged (F (6.176, 382.9) = 12.12; P<0.0001) and no effect of treatment on latency. n=17-18 rats per sex per treatment. ****P < 0.0001 C. Quantification of the latency to respond to gravitational cues. Repeated measures three-way ANOVA showed no effect of sex, data was then collapsed. Repeated measures two-way ANOVA showed decrease in latency to respond to gravitational cues as animals aged (F (4.896, 181.2) = 75.32; P<0.0001) and no effect of treatment on latency. n=17-8 rats per sex per treatment. ****P < 0.0001 D. Quantification of dam latency to retrieve pups. Repeated measures two-way ANOVA showed sex x treatment interaction (F (1, 35) = 4.882; P=0.0338). Bonferroni post hoc comparison between groups. n=9-11 rats per sex per treatment. *P < 0.05 E. Quantification of the latency to surface right. Repeated measures three-way ANOVA showed no effect of sex, data was then collapsed. Repeated measures two-way ANOVA showed decrease in latency to surface right as pups aged (F (2.027, 74.99) = 17.60; P<0.0001) and no effect of treatment on latency. n=17-18 rats per sex per treatment. ****P < 0.0001 F. Quantification of the latency to fall from a hanging wire. Repeated measures three-way ANOVA showed no effect of sex, data was then collapsed. Repeated measures two-way ANOVA showed increased latency to fall from a hanging wire as pups aged (F (3.544, 219.7) = 31.29; P<0.0001) and no effect of treatment on latency. n=17-18 rats per sex per treatment. ****P < 0.0001 G. Quantification of body weight, individual animal data points shown with mean line displayed due to overlapping averages. Mixed-effects analysis showed a main effect of sex (F (1, 60) = 246.9; P <0.0001) so analysis was repeated as two-way repeated measures ANOVAs to assess the effect of treatment separately in male and female rats. Mixed-effects analysis of females showed an increase in weight across age (F (1.782, 52.47) = 2676; P <0.0001) with no effect of treatment on weight. Mixed-effects analysis of males showed an increase in weight across age (F (1.442, 41.82) = 3142; P <0.0001) with no effect of treatment on weight. n=9-12 rats per sex per treatment. ****P < 0.0001 H. Schematic showing treatment paradigm, timeline, and legend for I-N. I. Quantification of open field center time. Two-way ANOVA showed no interaction or main effects of sex or treatment. n=11-14 rats per sex per treatment. J. Quantification of open field line crosses. Two-way ANOVA showed no interaction or main effects of sex or treatment. n=11-14 rats per sex per treatment. K. Quantification of recognition index during novel object recognition. Two-way ANOVA showed no interaction or main effects of sex or treatment. n=12-16 rats per sex per treatment. Horizontal dashed line indicates recognition index of 0.5 L. Quantification of the effect of postnatal THC on time spent in open arm of elevated plus maze. Two-way ANOVA showed no interaction or main effects of sex or treatment. n=15-17 rats per sex per treatment. M. Quantification of the total number of play events. Two-way ANOVA showed main effect of sex with males playing more than females (F (1, 59) = 4.424; P=0.0397) and a main effect of treatment with THC exposure increasing play (F (1, 59) = 4.807; P=0.0323). n=15-17 rats per sex per treatment. *P < 0.05 N. Quantification of the recognition index score. Two-way ANOVA showed main effect of treatment with THC exposure decreasing recognition index (F (1, 60) = 8.024; P=0.0063). **P < 0.01. One sample t-test against a hypothetical value of 0.5 of vehicle treated females (t=3.312, P=0.0047), THC treated males (t=1.485, P=0.1584), vehicle treated males (t=3.944, P=0.0015), and THC treated males (t=0.6330, P=0.5357). ##P < 0.01. n=15-17 rats per sex per treatment. Horizontal dashed line indicates recognition index of 0.5

Interestingly, while postnatal exposure to THC appeared to have no effect on development as well as many nonsocial juvenile behaviors, we found that it significantly altered juvenile social behaviors. Postnatal THC exposure increased the total number of play behaviors in the social play test (Figure 1M) and decreased the recognition index in the social recognition test (Figure 1N) in both female and male juvenile rats. Importantly, the recognition indices of THC-treated females and males were not significantly different than a hypothetical value of 0.5 (i.e. value that indicates equal time spent with the familiar and novel stimulus animals; one sample t-test), indicating that THC-treated rats had impaired social memory.

### Postnatal THC exposure alters microglial phagocytosis and newborn cell number in the developing amygdala

To assess whether the observed changes in juvenile social behaviors were correlated with altered microglial dynamics in the developing amygdala, we treated pups with both THC and bromodeoxyuridine (BrdU), to mark newborn cells, from P0-3 (Figure 2A). This time course and brain region were chosen based on our prior observation that endocannabinoid signaling directs microglia-mediated phagocytosis of newborn cells across the first 4-days of life in rats, which is responsible for establishing the sex difference in juvenile play behavior [13]. We analyzed the developing amygdala, determining its boundary using the optic tract (OT) (Figure 2B) at P4 and visualized microglia via immunohistochemistry using an antibody for ionized calcium binding adaptor molecule 1 (Iba1) (Figure 2D). Postnatal THC exposure did not alter the total number of microglia (Figure 2E), but increased the total number of phagocytic microglia, determined by the presence of a phagocytic cup (Figure 2C), in THC-treated females (Figure 2F). This was reflected by an increase in the percentage of phagocytic microglia in THC-treated females, with no effect in THC-treated males (Figure1G). We then quantified the number of newborn cells and found a corresponding decrease in BrdU+ cell number only in THC-treated females (Figure 1H). This being consistent with the notion that postnatal THC exposure increased phagocytosis in females, which lead to more newborn cells being phagocytosed, and a subsequent decrease in total newborn cell number.

**Figure 2:**
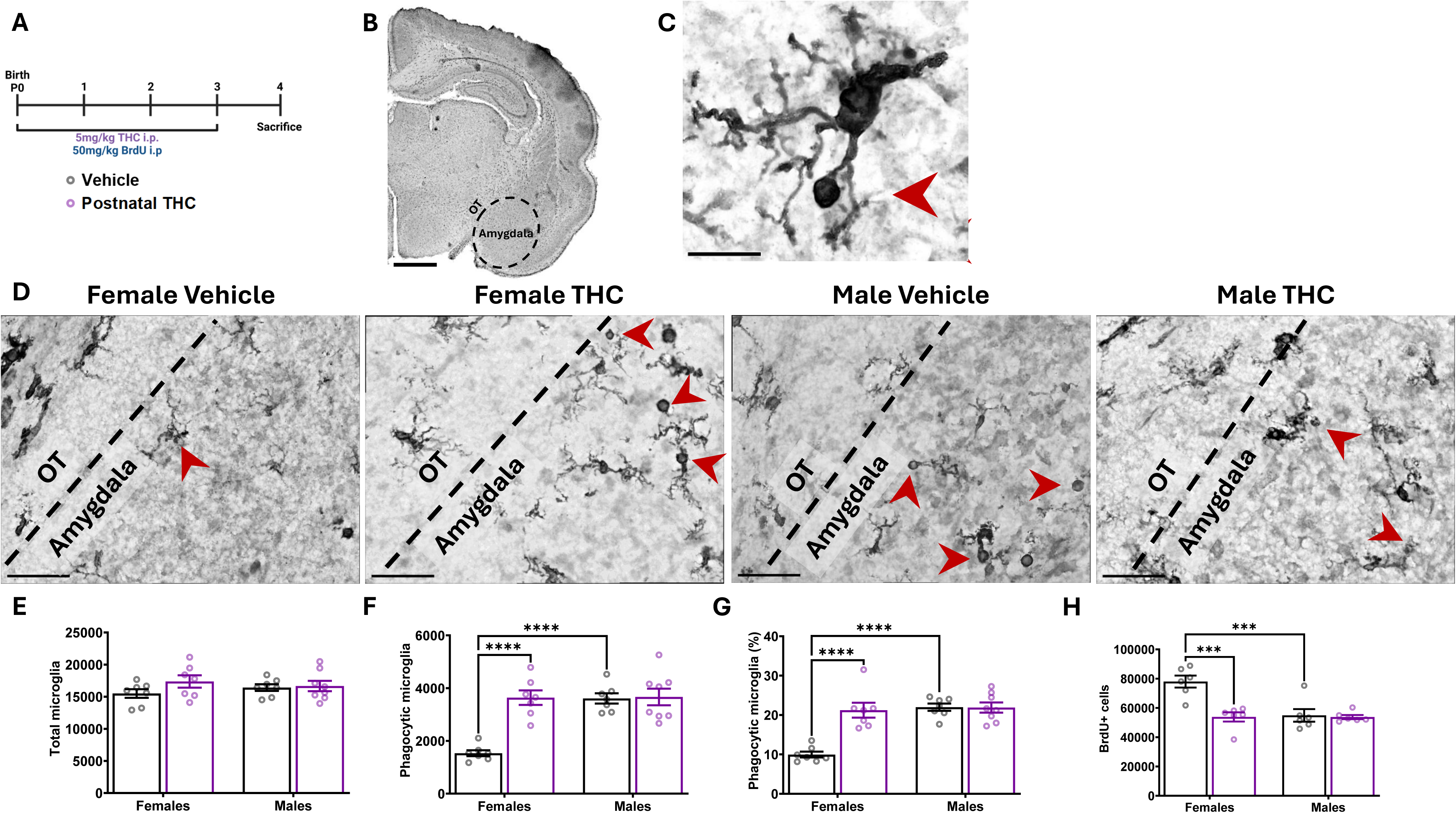
Postnatal exposure to THC alters microglial phagocytosis and newborn cell number in the developing amygdala. Bars represent the mean ± SEM, open circles represent individual data points for each animal. A. Schematic showing treatment paradigm, timeline, and legend for E-H. B. 4x Nissl and Iba1 stained coronal section of the P4 brain with dashed black line that indicates the boundaries of the developing amygdala used for analysis. Optic tract (OT) and amygdala denoted as landmarks for regions of analysis. Scale bar represents 1000um. C. 100x Nissl and Iba1 stained microglia denoting morphology of phagocytic cup (red arrow) used for quantification. Scale bar represents 15um. D. Representative 100x images of Nissl and Iba1 labeling from P4 developing amygdala. Black dashed line denotes boundary between amygdala and OT as landmark. Red arrowheads indicate phagocytic microglia. Scale bars represents 40 µm. E. Quantification of the total number of all microglia. Two-way ANOVA showed no interaction or main effects of sex or treatment. n=7-8 rats per sex per treatment. F. Quantification of the number of phagocytic microglia. Two-way ANOVA showed sex x treatment interaction (F (1, 25) = 17.42; P=0.0003). Bonferroni post hoc comparison between groups. n=7-8 rats per sex per treatment. ****P < 0.0001 G. Quantification of the percentage of microglia that are phagocytic. Two-way ANOVA showed sex x treatment interaction (F (1, 25) = 19.37; P=0.0002). Bonferroni post hoc comparison between groups. n=7-8 rats per sex per treatment. ****P < 0.0001 H. Quantification of the total number of BrdU+ cells in the developing amygdala. Two-way ANOVA showed sex x treatment interaction (F (1, 20) = 11.30; P=0.0031). Bonferroni post hoc comparison between groups. n=7-8 rats per sex per treatment. ***P < 0.001

### Prenatal exposure to THC alters social play and decreases social recognition without impacting development or other nonsocial behaviors

After observing altered social behavior and microglial dynamics following postnatal THC exposure, we sought to determine if these effects were specific to a defined developmental window. To test this, we exposed pups to THC *in utero* by treating pregnant dams with THC (5 mg/kg) from gestational day 15 (GD15) to GD21, then cross fostered pups at birth to an untreated lactating dam. We then assessed the development of the offspring using the same developmental behavior battery as before (Figure 3A). Similar to our findings above, prenatal THC exposure did not affect developmental locomotion, negative geotaxis reflex, dam latency to retrieve pups, surface righting reflex, wire hang ability, or body weight gain (Figure 3B-G). We again assessed juvenile behaviors in these same animals (Figure 3H) and found no changes in open field center time or line crosses, novel object recognition index, or open arm time in elevated plus maze (Figure 3I-L).

**Figure 3:**
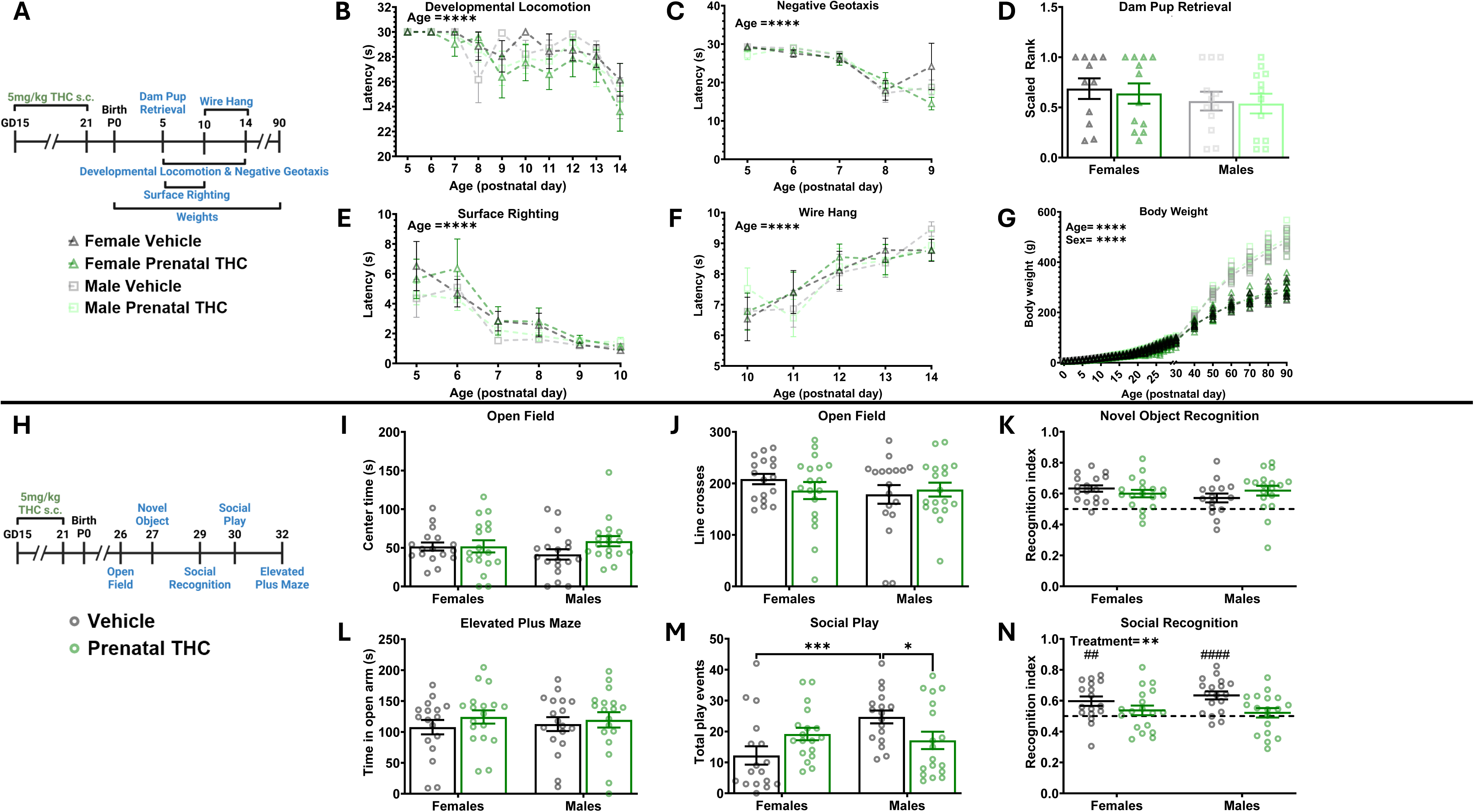
Prenatal exposure to THC alters social play and decreases social recognition while broadly not effecting development or other nonsocial behaviors. A. Schematic showing treatment paradigm, timeline, and legend for B-G B. Quantification of the latency to locomote. Repeated measures three-way ANOVA revealed no effect of sex, data was then collapsed. Repeated measures two-way ANOVA showed decrease in latency to locomote as animals aged (F (9, 621) = 8.404; P<0.0001) and no effect of treatment on latency. n=15-17 rats per sex per treatment. ****P < 0.0001 C. Quantification of the latency to respond to gravitational cues. Repeated measures three-way ANOVA revealed no effect of sex, data was then collapsed. Repeated measures two-way ANOVA showed decrease in latency to respond to gravitational cues as animals aged (F (3.925, 270.8) = 78.50; P<0.0001) and no effect of treatment on latency. n=9-11 rats per sex per treatment. ****P < 0.0001 D. Quantification of the dam latency to retrieve pups. Two-way ANOVA shoed no interaction or main effects of sex or treatment. n=11-12 rats per sex per treatment. E. Quantification of the latency to surface right. Repeated measures three-way ANOVA revealed no effect of sex, data was then collapsed. Repeated measures two-way ANOVA showed decrease in latency to surface right as animals aged F (2.354, 162.4) = 32.38; P<0.0001) and no effect of treatment on latency. n=9-11 rats per sex per treatment. ****P < 0.0001 F. Quantification of the latency to fall from a hanging wire. Repeated measures three-way ANOVA revealed no effect of sex, data was then collapsed. Repeated measures two-way ANOVA showed an increase in latency to fall from a hanging wire as animals aged (F (3.581, 247.1) = 14.51; P<0.0001) and no effect of treatment on latency. n=15-17 rats per sex per treatment. ****P < 0.0001 G. Quantification of body weight, individual animal data points shown with mean line displayed due to overlapping averages. Repeated measures three-way ANOVA showed a main effect of sex (F (1, 38) = 129.6; P<0.0001) so analysis was repeated as two-way repeated measures ANOVAs to assess the effect of treatment separately in male and female rats. Two-way ANOVA of females showed an increase in weight across age (F (1.515, 28.78) = 1759; P<0.0001) with no effect of treatment on weight. Two-way ANOVA of males showed an increase in weight across age (F (1.427, 27.11) = 3185; P <0.0001) with no effect of treatment on weight. n=11-17 rats per sex per treatment. ****P < 0.0001 H. Schematic showing treatment paradigm, timeline, and legend for I-N I. Quantification of open field center time. Two-way ANOVA showed no interaction or main effects of sex or treatment. n=17-18 rats per sex per treatment. J. Quantification of open field line crosses. Two-way ANOVA showed no interaction or main effects of sex or treatment. n=17-18 rats per sex per treatment. K. Quantification of recognition index during novel object recognition. Two-way ANOVA showed no interaction or main effects of sex or treatment. n=15-18 rats per sex per treatment. Horizontal dashed line indicates recognition index of 0.5 L. Quantification of the total number of play events. Two-way ANOVA showed sex x treatment interaction (F (1, 66) = 8.422; P=0.0050). Bonferroni post hoc comparison between groups. N=17-18 rats per sex per treatment. *P < 0.05, ***P < 0.001 M. Quantification of recognition index score. Two-way ANOVA showed main effect of treatment with THC exposure decreasing recognition index (F (1, 67) = 8.421; P=0.0050). **P <0.01. One sample t-test against a hypothetical value of 0.5 of vehicle treated females (t=3.173, p=0.0059), THC treated females (t=1.178, p=0.2549), vehicle treated males (t=5.159, p<0.0001), and THC treated males (t=0.7001, p=0.4933). ##P <0.01 ####P < 0.0001. n=15-17 rats per sex per treatment. Horizontal dashed line indicates recognition index of 0.5 N. Quantification of time spent in open arm of elevated plus maze. Two-way ANOVA showed no interaction or main effects of sex or treatment. n=17-18 rats per sex per treatment.

In line with our prediction that the timing of THC exposure would change its impact on behavior, we found that prenatal THC exposure decreased the total number of play behaviors in males, while females were unaffected. (Figure 3M). Prenatal THC exposure also decreased the social recognition index in both sexes (Figure 3N). For both females and males, the recognition index in THC-exposed groups was not statistically different than a hypothetical value of 0.5 while the vehicle treated animal’s index scores were, once again indicating an impairment in social memory.

### Prenatal exposure to THC decreases microglial phagocytosis and newborn cell number in the developing amygdala

Because prenatal exposure to THC had distinct effects on social behavior when compared to postnatal exposure to THC, we next sought to determine if microglia were also differentially altered in the developing amygdala. To test this, we again exposed pups to THC *in utero* by treating pregnant dams from GD15-21, cross fostered offspring at birth, and subsequently treated pups with BrdU from P0-P3 to label newborn cells (Figure 4A). We quantified microglia (Figure 4B) and found that prenatal exposure to THC surprisingly decreased the total number of microglia in both sexes (Figure 4C) and decreased the number of phagocytic microglia (Figure 4D). However, the decrease in total and phagocytic population of microglia was proportional, meaning that the percentage of phagocytic microglia was unchanged (Figure 4E). Interestingly, the decrease in phagocytosis did not correspond to an increase in the number of newborn cells in the developing amygdala; rather, we found that prenatal THC exposure also reduced BrdU+ cell number in both sexes (Figure 4F).

**Figure 4:**
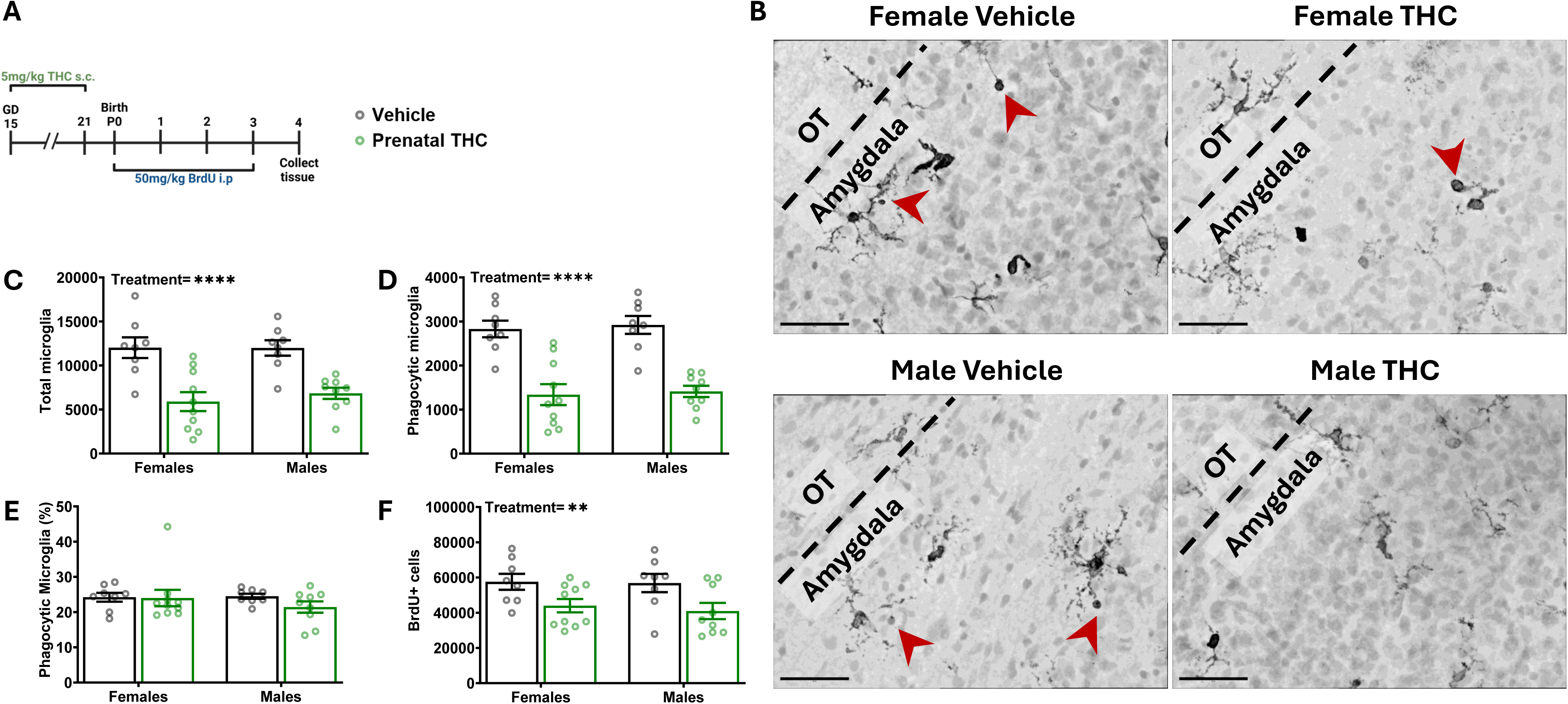
Prenatal exposure to THC diminishes microglial population transiently during development. A. Schematic showing treatment paradigm, timeline, and legend for C-F. B. Representative 100x images of Nissl and Iba1 labeling from P4 developing amygdala. Black dashed line denotes boundary between amygdala and OT as landmark. Red arrowheads indicate phagocytic microglia. Scale bars represent 40 µm. C. Quantification of the total number of all microglia. Two-way ANOVA showed main effect of treatment with THC exposure decreasing the number of microglia (F (1, 31) = 33.76; P<0.0001). n= 8-10 rats per sex per treatment. ****P < 0.0001 D. Quantification of the number of phagocytic microglia. Two-way ANOVA showed main effect of treatment with THC exposure decreasing the number of phagocytic microglia (F (1, 31) = 57.16; P<0.0001) n= 8-10 rats per sex per treatment. ****P < 0.0001 E. Quantification of the percentage of microglia that are phagocytic. Two-way ANOVA showed no interaction or main effects of sex or treatment. n= 8-10 rats per sex per treatment. F. Quantification of the total number of BrdU+ cells. Two-way ANOVA showed main effect of treatment with prenatal THC decreasing the total number of BrdU+ cells (F (1, 31) = 10.65; P=0.0027). n= 8-10 rats per sex per treatment. **P < 0.01

### Prenatal exposure to THC diminishes microglial population transiently during development

To further investigate the decrease in microglia number at P4 following prenatal THC exposure, we quantified the microglia proliferation within the developing amygdala. We treated pregnant dams with THC from GD15-21 to expose pups *in utero*, then treated pups with BrdU on P0 to label newborn cells, followed by euthanasia 3 hours post injection (Figure 5A). We then quantified the number of microglia, as well as proliferating microglia (BrdU+) in the developing amygdala (Figure 5B). Paradoxically, we found no differences in microglia number at P0 (Figure 5C) and found an increase in BrdU+ microglia (Figure 5D). We interpret these findings to suggest that the decrease in microglia seen at P4 occurs after the cessation of THC exposure between P0 and P4, rather than being an acute effect of prenatal THC exposure.

**Figure 5:**
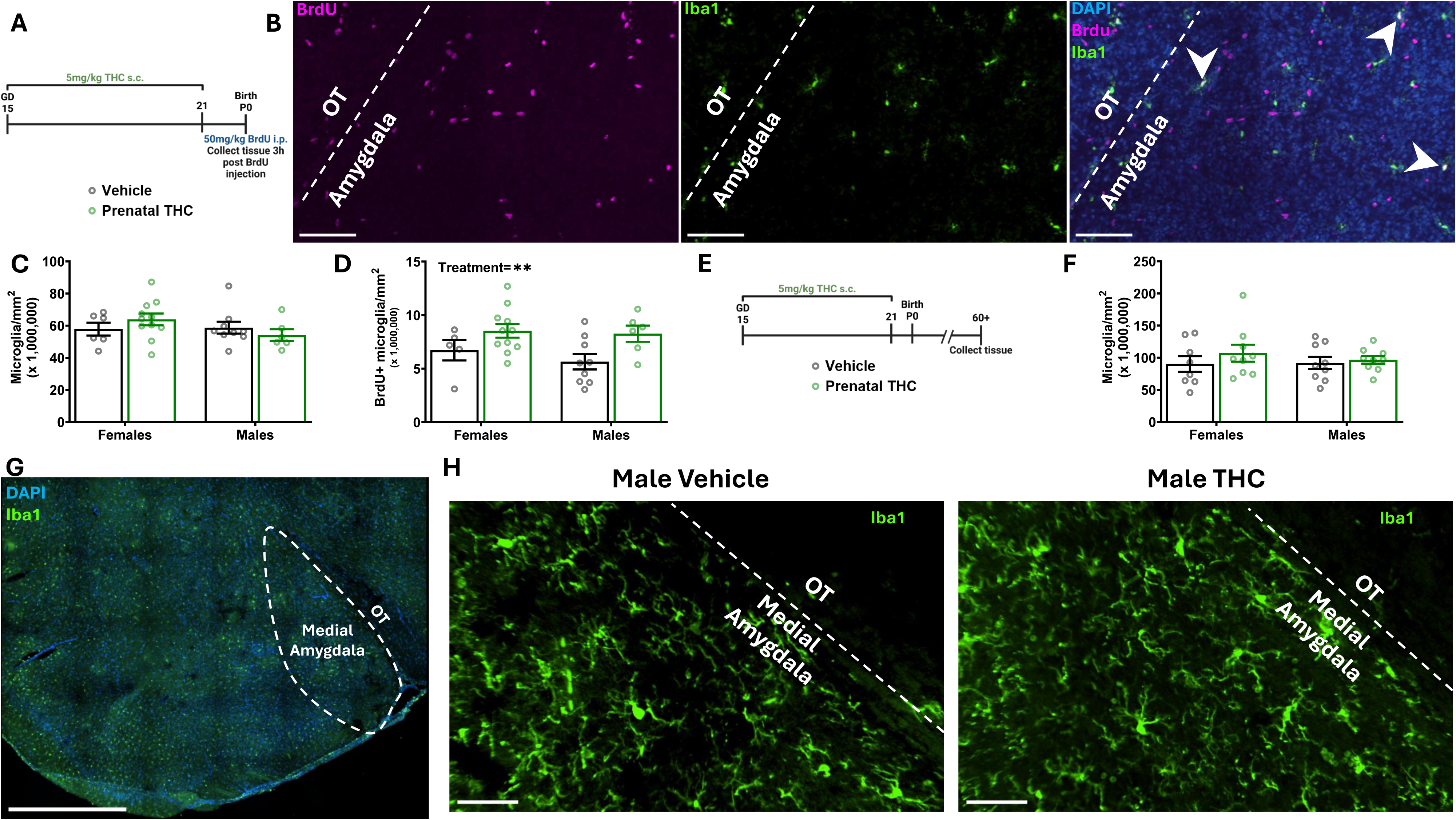
Prenatal exposure to THC alters microglial timeline temporarily. Bars represent the mean ± SEM. Open circles represent individual data points for each animal A. Schematic showing treatment paradigm and timeline for C-D. B. Representative 20x image of Iba1, DAPI, and BrdU labeling from P0 developing amygdala. Separate images with BrdU and Iba1 channels individually for clearer viewing. White dashed line denotes boundary between amygdala and OT as landmark. White arrowheads indicate proliferating microglia (double positive for Iba1 and BrdU). Scale bars represent 100µm. C. Quantification of the density of microglia. Two-way ANOVA showed no interaction or main effects of sex or treatment. n=6-11 rats per sex per treatment. D. Quantification of the density of Brdu+ Microglia. Two-way ANOVA showed main effect of treatment with THC exposure increasing the density of BrdU+ microglia (F (1, 27) = 7.889; P=0.0091) n=6-11 rats per sex per treatment. **P < 0.01 E. Schematic showing treatment paradigm and timeline for F F. Quantification of density of microglia in adulthood. Two-way ANOVA showed no interaction or main effects of sex or treatment. n=8-9 rats per sex per treatment. G. Representative 20x image of Iba1 and DAPI from P60+ amygdala. Dashed line denotes medial amygdala. Scale bar represents 1000µm. H. Representative 20x images of Iba1 from P60+ amygdala. Vehicle and THC treated animals to show microglial populations. White dashed line denotes boundary between amygdala and OT as landmark. Scale bars represent 50µm.

Finally, to determine whether the changes in microglia number after prenatal THC exposure endured into adulthood, we treated pregnant dams with THC from GD15-21 and subsequently allowed the offspring to grow to adulthood (P60+) (Figure 5E). We then quantified microglia within the medial amygdala (Figure 5G-H) and found no differences in the number of microglia in either sex (Figure 5F), indicating that the decrement in microglia number is temporary during development.

## Discussion

The primary psychoactive component of cannabis, THC, is a potent modulator of the eCB system’s cannabinoid receptors [9]. Cannabis use among pregnant populations is relatively understudied and the need for models of developmental exposure is evident when considering that the amount and timing of exposure to substances such as alcohol during pregnancy can result in vastly different outcomes [6–8]. It is possible that differences reported in cognitive and behavioral outcomes of children exposed to THC [28] could be a consequence of the developmental period during which exposure occurred. Here using a rodent model we find that the timing of THC exposure resulted in some common and some distinct changes to juvenile social behavior while leaving many developmental milestones and behaviors unchanged. Specifically with regards to social play, the changes in play behavior were dependent upon the timing of THC exposure and were accompanied by distinct effects on microglial dynamics within the developing amygdala.

### THC exposure during the prenatal and postnatal periods in rats does not affect the acquisition of normal developmental milestones and non-social behaviors

We found that neither exposure paradigm (prenatal or postnatal) altered the acquisition of major developmental milestones, and did not affect juvenile locomotor, object memory, or anxiety-like behaviors (Supplemental Table 1). While in humans prenatal THC exposure has been associated with increased anxiety and anhedonia in adolescence [29], other groups working on rodent models have reported no effect on anxiogenic behavior [30–32]. Additionally, studies of prenatal THC exposure in rodents often report mixed effects on birth weight [33, 34], locomotion [33, 35], and ultrasonic vocalizations (USVs) [36, 37], with a meta-analysis suggesting low weighted effect size of THC exposure on behavior and locomotor activity [38]. Here we find only postnatal THC exposure disrupted dam-pup interactions: dams retrieved control male pups faster than females and postnatal THC exposure equalized latencies, delaying male pup retrieval. This change may stem from alterations in ultrasonic vocalizations (USVs), as male pups vocalize more frequently than females which promotes quicker retrieval by the dam [39]. Other reports show that prenatal THC exposure increases pup USVs [40], however, prenatal THC exposure did not alter pup retrieval suggesting that pup-dam communication is unaffected in our paradigm. Overall, these findings indicate THC exposure during these distinct time periods does not dramatically alter development or impact non-social behaviors by the juvenile age.

### Juvenile social behavior is specifically altered based on timing of THC exposure

Social play is one of the earliest pro-social behaviors, is highly conserved, and is most robustly displayed during the juvenile period [41–43]. In nearly every species that plays, males engage in more frequent and vigorous rough-and-tumble play than females [44]. The amygdala controls sex differences seen in social play [14], and we have previously shown this difference in rats to be mediated by eCB signaling in the developing amygdala [13]. Given that THC is a potent activator of eCB receptors, we hypothesized THC exposure might alter the expression of juvenile play behavior. We found that the effects of THC on later-life play behavior were specific to the time of exposure: prenatal THC exposure reduced play behavior in males but did not affect females, while postnatal THC exposure increased play behavior in both sexes.

Both THC exposure paradigms resulted in impaired social memory in both sexes, highlighting the behavioral specificity of the effects of THC exposure. These alterations in social behaviors are consistent with similar work showing cannabinoid exposure altering social behaviors [30, 40, 45, 46]. Given that we found no effect of either THC exposure paradigm on locomotor or anxiety-like behaviors, it seems likely that the observed changes in play are due to specific changes in the development of social behavior circuitry [47, 48]. Furthermore, we found no changes in novel object recognition memory following either THC exposure paradigm, suggesting that specifically social memory, not memory overall, was impaired. This points to both a time dependent and region dependent effect of THC exposure, which could be explained by the timing of neurogenesis and development of various brain structures during early development [15, 49].

A further caveat to the current findings is the still limited understanding of the nature of sex differences in social play which is scored based on the frequency of pre-determined “elements” such as pouncing, pinning and boxing. The advent of machine learning tools to use unbiased metrics to observe the patterns and syllables of play may reveal heretofore unknown complexity in play elements [50, 51], and that shifts in the nature of play rather than just the amount could be occurring due to THC exposure. This warrants further characterization to how the quality of play could also be changed following THC exposure at varying timepoints in both sexes.

### Microglial dynamics in the developing amygdala are specifically altered based on timing of THC exposure

Microglia are critical for neurodevelopment, through production of neurotrophic factors and cytokine secretion, synapse pruning, and, relevant to our study, phagocytosis. [52–54]. We found that postnatal THC exposure increased microglial phagocytosis in females but not males and was accompanied by a reduction in BrdU+ cells, consistent with our previous findings demonstrating engulfment of astrocyte progenitors in the developing amygdala. Increased phagocytic activity, and therefore fewer astrocyte progenitors in females, may explain the observed increase in social play, as a lower astrocyte density within the medial amygdala is permissive for higher levels of play [13]. In males, however, postnatal THC exposure increased social play without detectable changes in phagocytosis, suggesting THC exposure may alter the development of social behavior circuitry independent of microglial dynamics. These effects could be related to findings showing that cannabinoid exposure can delay the GABA switch [39], ablate eCB mediated long-term depression in the prefrontal cortex as well as nucleus accumbens [30, 45], alter dopaminergic activity [55], and dysregulate striatal epigenetic expression [56].

How eCBs directly influence phagocytic state is unknown, but postnatal THC exposure could be inducing this state directly or it could be indirectly modulating the eCB system by changing the levels of circulating eCBs or their synthetic and degrative enzymes. The eCB ligand 2-AG has been implicated to be a chemoattractant “find me” signal for microglia [57] and thus by modulating its synthetic or degrative enzymes, THC exposure could be indirectly increasing this “find me” signal. Activation of CB1 during adolescence has been shown to mediate microglial apoptosis [58], while activation of CB2 has been shown to not only switch microglia to their more anti-inflammatory phenotype [59], but also switch microglia to their more activated phagocytic phenotype [60]. These highly variable outcomes emphasize the importance of investigating the specific microglial profiles following THC exposure at specific timepoints to determine how alterations in their function may impact brain development.

Prenatal THC exposure reduced the microglia population and microglial phagocytosis in both sexes during the postnatal period. Paradoxically, the decrease in phagocytosis did not increase, but rather decreased newborn cell number in both sexes following prenatal THC exposure. Microglia secrete important trophic factors that promote newborn cell proliferation and survival, namely tumor necrosis factor alpha (TNFα) [61], and THC exposure may alter the expression of these factors during development. THC has been shown to be potently anti-inflammatory by decreasing MIP-1α and MIP-1β in vitro [62], as well as reducing the levels of IL-6, IL-8, and TNF-α in vitro [63]. However, its exact mechanism of action during development remains understudied.

After cessation of prenatal THC treatment, the microglia population rebounds rapidly via microglia proliferation postnatally. This suggests that microglia during prenatal development may be uniquely sensitive to the effects of THC, and the functional consequences of in utero exposure may be dramatically different than postnatal exposure in rodents. Prenatal THC exposure could be altering the distinct transcriptomic temporal stages of microglial developmental [64]. Similarly, given THC’s effect on chemotaxis [65, 66], the normal colonization and migration patterns of microglia may be interrupted during development.

In conclusion, our findings demonstrate the importance of modeling different periods of gestational cannabis exposure by showing that the timing of THC exposure has differential effects on social behavior as well as microglial dynamics during development. Interestingly, our findings show that social behaviors are uniquely sensitive to disruption by THC exposure, whereas major developmental milestones and behaviors associated with cognition and emotional reactivity were unaffected. Play behavior is critical for the development of later-life social behaviors [67–69]; thus, the behavioral changes due to THC exposure in juvenile animals may have lasting impacts on behavior during adulthood. As society decreases stigma surrounding cannabis, and increases access, it will be increasingly important to understand the enduring but latent effects of use during pregnancy.

## Data Availability Statement

The datasets generated during and or/analyzed during the current study are available from the corresponding author on reasonable request.

## Author Contributions

Conceptualization, JVR, AEM, MMM; Methodology, JVR, AEM, MMM; Formal Analysis, ALP, JVR; Investigation, ALP, AEM, KNS, KRM, JVR; Writing-Original Draft, ALP, JVR, MMM; Writing-Review and Editing, ALP, JVR, MMM; Supervision, JVR, MMM; Funding Acquisition, MMM.

## Funding

This work was supported by R01 5R01DA039062-08 to MMM

## Competing Interests

The authors have nothing to disclose.

## Notes

### Competing Interest Statement

The authors have declared no competing interest.

