## Supplemental Methods for "Timing matters: modeling the effects of gestational cannabis exposure on social behavior and microglia in the developing amygdala"

Immunohistochemistry

For fluorescent imaging, slide-mounted sections were washed in PBS and blocked with either 10% bovine serum albumin (BSA) in PBS + 0.4% Triton X-100 (PBS-T) for 1 h. Slides were incubated in primary antibody solution (5% BSA in PBS-T) overnight. The following day, slides were incubated in secondary antibody solution (5% BSA in PBS-T) for 2 h and coverslipped with ProLong Diamond Antifade (Thermo Fisher Scientific).

For DAB staining, slide-mounted sections were washed in Tris-buffered saline (TBS; 0.05M, pH 7.6), incubated in 0.3% hydrogen peroxide in TBS for 30 min at room temperature. Sections were blocked with 10% BSA in TBS + 0.4% Triton X-100 (TBS-T). Sections were incubated in primary antibody in solution (5% BSA in TBS-T) overnight. The following day, sections were incubated in biotinylated secondary antibody for 1 h, followed by incubation in ABC reagent (1:500 dilution; Vectastain Elite ABC Kit, Vector Laboratories) in TBS-T for 1 h, and visualized using DAB chromagen or nickel-enhanced DAB chromagen (0.05% 3,3′-diaminobenzidine, 0.2% nickel (II) sulfate, 0.006% hydrogen peroxide; all from Sigma-Aldrich). The DAB reaction was allowed to proceed until completion, as confirmed under a microscope. Sections were counterstained with either hematoxylin or methyl green, dehydrated in an ascending ethanol series, defatted in xylene, and coverslipped with DPX mounting medium.

Primary antibodies used included the following: Rabbit anti-Iba1 (1:1000; Wako Cat#019-19741, RRID:AB_839504), mouse anti-BrdU (1:500; BD Biosciences Cat#347580, RRID:AB_400326), Secondary antibodies used included the following: biotinylated anti-rabbit (1:500; Vector Laboratories Cat#BA-1000, RRID:AB_2313606), biotinylated anti-mouse (1:500; Vector Laboratories Cat#BA-2000, RRID:AB_2313581), Alexa Fluor 488- or 594-conjugated antibodies to rabbit (488 Cat#A21206, RRID:AB_2535792; 594 Cat#A21207, RRID:AB_141637), or mouse (488 Cat#A21202, RRID:AB_141607; 594 Cat#A21203, RRID:AB_2535789).

Unbiased stereological cell counting

Every third section (45µm thick) was used for analysis for a total of four sections, and both hemispheres of the amygdala were quantified. The boundaries of each region were drawn with a 4x objective, referencing a neonatal rat atlas [70], and the optical fractionator method was used to quantify microglia and BrdU+ cells at 40x magnification, using a 100µm x 100µm counting grid with a 250µm x 250µm sampling grid for microglia and a 50µm x 50µm counting grid with a 250µm x 250µm sampling grid for BrdU+ cells. The optical dissector height was set to 12µm with a 2µm guard zone on the top and bottom for both quantifications. Microglia were counted based on the presence of an observable cell body within the designated counting region and determined to be phagocytic if the microglia contained an observable phagocytic cup that was distinctly identifiable from the cell body. BrdU+ cells were counted if the nuclear staining was uniformly dark and was within the designated counting region.

Quantification of cell density

The surface tool was used to create volumetric boundary of either the developing amygdala for neonatal studies, or the medial amygdala in adults. To determine the density of microglia within these regions, individual microglia were counted using the automatic spots function which counted microglia based on their Iba1 positive soma and cell size of approximately 13µm. Density was then calculated by dividing the number of microglia by the volumetric area of the determined region. Proliferating microglial density was determined by using the spots function to automatically count cells that were double positive for both Iba1 and BrdU, and then dividing that total by the area of the volumetric boundary.

Developmental battery

*Weight:* Weights of animals were taken daily from P0-90.

*Dam Latency to Retrieve Pups:* On P5 each pup was placed one at a time opposite the nest and the time it took the dam to retrieve the pup was measured. Data was rank ordered within litter with the latency being assigned a value 1-12, with the rank then being divided by the total litter size to scale the data with the fastest being 1/12 and the slowest being 12/12.

*Surface Righting:* From P5-10 each pup was placed on its back on a flat surface. The time for the pup to return to its four limbs was recorded with a cutoff time of 30 seconds.

*Developmental Locomotion:* From P5-15 each pup was placed on a flat surface in the center of an underlaid circle 13cm in diameter. The time for all four legs of the pup to exit the circle was recorded with a cutoff time of 30 seconds.

*Negative Geotaxis:* From P5-15 each pup was placed face down on a ramp with a 25-degree incline that was covered with a wire mesh to enable traction. The time for the pup to turn 180 degrees with its head and trunk oriented to the top of the incline was recorded with a cutoff time of 30 seconds.

*Wire Hang:* From P10-15 each pup's forepaws were placed against a horizontal wire rod suspended above a padded bin. The duration of time each pup hung from the wire was recorded with a cutoff time of 10 seconds.

Juvenile behavioral battery

The behavioral battery was performed as previously described [26]. All behavior testing took place during the dark phase of the animal's light/dark cycle under red light illumination. Animals were weaned on P21 and housed in same-sex, same-treatment sibling pairs. Details for each behavior can be found in the supplementary methods.

*Open Field:* On P26 animals were tested for 10 min in an open field (78 × 78 cm, 40 cm high), underlaid with a grid delineating perimeter and center regions. Center time and line crosses were recorded.

*Novel Object Recognition:* On P27 animals were placed in the same arena used for open field testing and were exposed to a pair of identical objects placed on opposite ends of the arena for 5 min. After this exposure, animals were placed back in their home cages for 1 h and then were returned to the arena and exposed to a now familiar object as well as a novel object. The time spent investigating each object was recorded, and the recognition index was calculated (time with novel object / time with novel object + time with familiar object). The position of objects in the arena and the order of object exposure was counterbalanced between treatment groups.

*Social Recognition:* On P29 animals were singly housed in a test cage identical to their home cage with ad libitum access to food and water for 2 h to allow for habituation. A sex and age matched stimulus rat was then placed into the cage with the test rat for 4 min and the number of time the experimental animal spent interacting with the stimulus animal was recorded. The stimulus rat was then removed, and the test rat remained in the test cage for a retention interval of 20 min. The original familiar stimulus rat and a novel stimulus rat were then placed into the test cage with the test rat. The time the test rat spent investigating each of the stimulus rats was recorded and recognition index was calculated (time with novel rat / time with novel rat + time with familiar rat).

*Social Play:* Protocol was performed as described [71]. On P30 animals were isolated for two hours and then same-sex, same-treatment non-sibling pairs of animals were placed in an enclosure (49 × 37 cm, 24 cm high) with TEK-Fresh cellulose bedding. Animals were allowed to acclimate for 2 min, then video recorded for 10 min. Videos were then scored offline to determine the number of pounces, pins, and boxing behaviors to measure social play.

*Elevated Plus Maze:* On P32 animals were individually placed on an elevated plus-shaped polycarbonate apparatus with two open and two enclosed arms and video recorded for 5 minutes. The time spent by each animal in the open arms compared to the closed arms was then used as a measure of anxiety-like behavior.

Quantification and statistical analysis

To assess possible litter effects for all studies that were performed during gestational timepoints, data was initially analyzed *a priori* using the dam as the n (i.e. response of all pups in a litter averaged). No differences were found between significant data whether the dam was the n or whether individual offspring were the n, indicating no significant litter effects in our experiments. Data including multiple timepoints were analyze *a priori* using a three-way analysis of variance (ANOVA) to determine if there was a main effect of sex, and when no main effect of sex was found, sex was collapsed, and data were then analyzed using a two-way ANOVA for age and treatment. Data including multiple experimental groups were analyzed using two-way ANOVA for treatment and sex when appropriate. A one sample t-test against a theoretical value of 0.5 to determine if recognition differed from random chance was used.
