## Supplemental Table 1 for "Timing matters: modeling the effects of gestational cannabis exposure on social behavior and microglia in the developing amygdala"

*Supplemental Methods:*

*Supplemental Table 1: Summary table of behavioral effects of THC treatment paradigms*

Summary table showing the behavioral assays performed, the age they were performed at, and the effects either postnatal or prenatal THC had on them for both the developmental battery as well as the battery performed during the juvenile period.

| Behavior | Age | Postnatal THC (P0-7) | Prenatal THC (GD15-21) |
|---|---|---|---|
| **Developmental Battery** | | | |
| **Body Weight** | P0-90 | No Effect | No Effect |
| **Dam Latency to Retrieve Pups** | P5 | **Decreased latency to retrieve females**<br>**Increased latency to retrieve males** * | No Effect |
| **Surface Righting** | P5-10 | No Effect | No Effect |
| **Developmental Locomotion** | P5-15 | No Effect | No Effect |
| **Negative Geotaxis** | P10-15 | No Effect | No Effect |
| **Wire Hang** | P10-15 | No Effect | No Effect |
| **Behavioral Battery** | | | |
| **Open Field; Center Time** | P26 | No Effect | No Effect |
| **Open Field; Line Crosses** | P26 | No Effect | No Effect |
| **Novel Object Recognition** | P27 | No Effect | No Effect |
| **Social Recognition** | P29 | **Decreased social recognition in females**<br>**Decreased social recognition in males** * | **Decreased social recognition in females**<br>**Decreased social recognition in male** * |
| **Social Play** | P30 | **Increased social play in females**<br>**Increased social play in males** * | **No effect in females**<br>**Decreased social play in males** * |
| **Elevated Plus Maze** | P32 | No Effect | No Effect |
